# Does AlphaFold predict the spatial structure of a protein from physics or recognize it (its main parts and their association) using databases?

**DOI:** 10.1101/2022.11.21.517308

**Authors:** Alexei V. Finkelstein

**Affiliations:** Institute of Protein Research, Russian Academy of Sciences, Pushchino, Moscow Region, Russian Federation

**Keywords:** bioinformatics, similarity of 3D structures, random sequence similarity, sequence identity, databases

## Abstract

The great success of the AlphaFold programs poses the following questions: (i) What is the main reason for this success? (ii) What exactly do AlphaFolds do: *prediction* of the 3D protein structure based on its amino acid sequence and knowledge of the protein physics or *recognition* of this 3D structure, based on the similarity between some parts of its amino acid sequence and parts of sequences with already known 3D structures? The answers given in this paper are: The main reason for the tremendous success of the AlphaFold is (i) the usage of huge protein databases, which already cover all or almost all of the protein superfamilies existing in nature; (ii) using these databases and the resulting multiple sequence alignments and coevolutionary information (like correlations in pairs and especially in triplets of amino acid residues in the contacting chain regions), AlphaFold *recognizes* a 3D structure of the examined amino acid sequence by a similarity of this sequence (or its parts) to related sequences with already known 3D structures. Concluding, I have to emphasize that this paper does not diminish the merit and utility of AlphaFold; it only explains the basis of its success.

## 1. Introduction

The great success of The AlphaFold and then AlphaFold 2 programs [1, 2] in identifying 3D (three-dimensional) protein structures from their amino acid (a.a.) sequences and the subsequent application of the latter program to the 3D structures of huge protein machines [3] raised two important questions: (i) What is the main reason for this success? (ii) What exactly do AlphaFolds do: *prediction* of the 3D protein structure based on its a.a. sequence and knowledge of the protein chain physics or *recognition* of this 3D structure based on the similarity between some parts of its a.a. sequence and parts of sequences with already known 3D structures?

Our approach to the latter dilemma will exploit a well-known (in bioinformatics) observation [4–7] that 20-25% or more identity of a.a. sequences is, as a rule, sufficient to ensure a high similarity of 3D protein structures (with a small, ≈2Å or less, difference between them). More specifically [5, 6], the residue identity below ≈20% in the pairwise alignments of sequences usually does not provide the correct alignment of 3D structures; the residue identity of 20-25% corresponds to the zone where the alignments of sequences may correspond, but also may be rather different from the alignments of 3D structures; and only the residue identity above ≈25% ensures coincidence of alignments of 3D protein structures and sequences.

## 2. Results

### 2.1. Theory

Now, let us estimate the expected similarity of a randomly taken a.a. sequence fragment *S_n_* (of ***n*** a.a. residues) to the closest in similarity chain fragment *S*’ from a set **Σ**_*N*_ of ***N*** other random sequences of the same size. In fact, we pose a question as to whether the set **Σ**_*N*_ is large enough to contain a sequence *S*’, whose high sequence identity to *S_n_* (over 20-25%) ensures high similarity of 3D structures of *S* ‘ and *S_n_*.

According to the Poisson distribution, the probability that the random a.a. sequence *S_n_* matches another random a.a. sequence of the same length ***n*** in ***m*** positions is

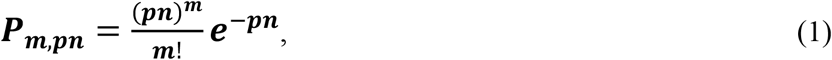

when each of the amino acids falls out with probability ***p***.

If the random sequence *S_n_* is compared not with one but with ***N*** random sequences (forming the set **Σ**_***N***_ of sequences the same length ***n***), the expected number of the set ***Σ***_***N***_ members matching *S_n_* in m positions is ***P_M,pn_N***. Thus, the equation ***P_M,pn_N*** = 1 determines the maximal expected number ***M*** of matches of the sequence *S_n_* with the most similar to it sequence from the set **Σ**_***N***_ of the ***n***-residue random sequences.

Given ***p*** << 1 (for proteins, ***p***≈1/20), not too small math expectation of the number of randomly matching amino acids (1 << ***pn***), long enough sequences (***pn*** << ***m***), and not too small actual sequence identity of similar sequences (1 << ***pn*** << ***m***), one can use the Stirling’s approximation [***m***! ≈ (***m***/ ***e***)^***m***^, where ***e***≈2.72] for factorials and get

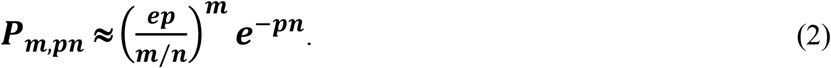

Thus, the value of ***M***/***n*** (where ***M*** is the maximal, in the “best” sequence *S*’ from the set **Σ**_***N***_, expected number of matches ***m*** with the given sequence *S_n_*) can be found from the following equation: 1/***P_M,np_*** = ***N***, or

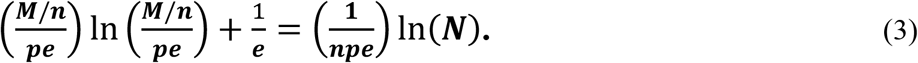

### 2.2. Numerical estimates

For ***p*≈1/20** (typical of proteins), ***n***≈**100** (typical protein domain size) and ***N***≈1.5×10^5^ (the number of spatial protein structures in the Protein Data Bank (PDB) in 2020, https://www.rcsb.org/stats/growth/growth-protein), ***M/n*** ≈ 0.19 (see Table 1). For ***p*≈1/20**, ***n*≈100** and ***N***≈1.9×10^8^ (the number of protein sequences in the UniProtKB database in 2020, https://academic.oup.com/nar/article/49/D1/D480/6006196), ***M/n*** ≈ 0.23 (see Table 1).. Thus, the search for the highest similarity between a random 100 a.a.-long sequence and the about 1.5×10^5^ PDB-stored protein chain sequences is expected to discover an approximately 19%-identical sequence; while a 23%-identical chain should be discovered among the about 1.9×10^8^ UniProtKB-stored protein sequences (see the blue bar in Fig. 1). Such a sequence identity level is usually sufficient to ensure close (with RMSD≈1.7±0.5Å, see Ref. 4) similarity of 3D structures of the examined protein to its closest analog in the databases of 2020.

**Figure 1.**
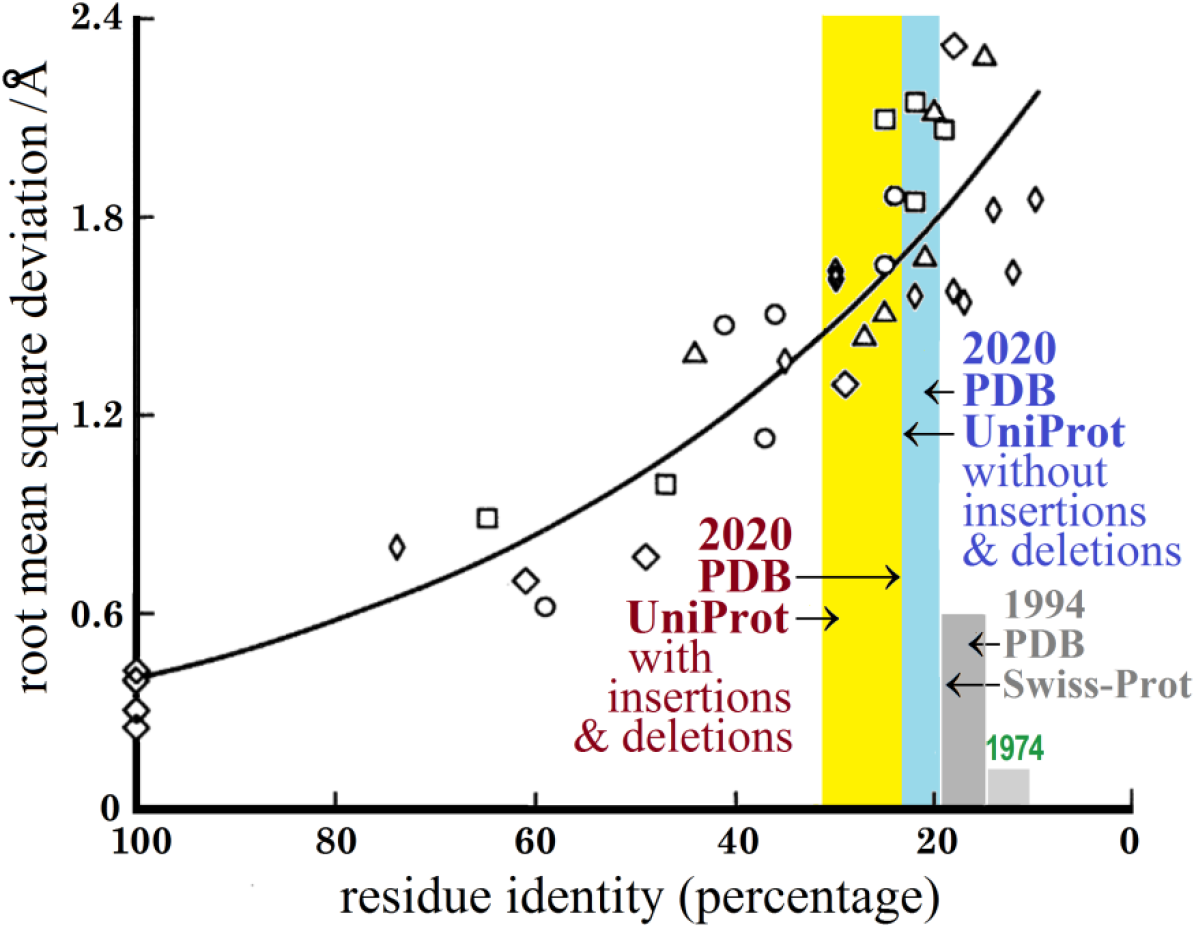
Structural divergence of homologous proteins plotted against the sequence identity (black symbols and curve, adapted from Ref. 4), and the expected ranges (colored bars, see explanations in the text) of residue identity for a “domain-size” (***n***≈100) random sequence to the most similar to it chain from UniProtKB and PDB databases of different years. The structural difference is measured by the root mean square deviations (Å) of the main-chain atomic positions of residues of the “protein cores” (comprising the main secondary structure elements and covering >90% of the chains with a >50% residue identity, and ≈50% of the chains with a ≈20% residue identity) [4] that have been optimally superimposed. The sequence similarity is measured in the percentage of residues that are identical in the superimposed cores. Different black symbols refer to different protein families.

**Table 1.** The ***M/n*** values obtained from the solution of equation (3) at different values of ***N*** and ***n***

| $n$ | $p$ | $N$ | $M/n$ |
| --- | --- | --- | --- |
| any | any | 1 | $p$ |
| 50 | 0.05 | 10 | $0.97 \times pe = 0.13$ |
| 50 | 0.05 | 1000 | $1.53 \times pe = 0.21$ |
| 50 | 0.05 | $1.5 \times 10^5$ | $2.00 \times pe = 0.27$ |
| 50 | 0.05 | $1.9 \times 10^8$ | $2.41 \times pe = 0.33$ |
| 50 | 0.05 | $2 \times 10^{14}$ | $3.39 \times pe = 0.46$ |
| 100 | 0.05 | 10 | $0.77 \times pe = 0.10$ |
| 100 | 0.05 | 1000 | $1.13 \times pe = 0.15$ |
| 100 | 0.05 | $1.5 \times 10^5$ | $1.43 \times pe = 0.19$ |
| 100 | 0.05 | $1.9 \times 10^8$ | $1.68 \times pe = 0.23$ |
| 100 | 0.05 | $2 \times 10^{14}$ | $2.28 \times pe = 0.31$ |
| 200 | 0.05 | 10 | $0.64 \times pe = 0.09$ |
| 200 | 0.05 | 1000 | $0.88 \times pe = 0.12$ |
| 200 | 0.05 | $1.5 \times 10^5$ | $1.07 \times pe = 0.15$ |
| 200 | 0.05 | $1.9 \times 10^8$ | $1.23 \times pe = 0.17$ |
| 200 | 0.05 | $2 \times 10^{14}$ | $1.61 \times pe = 0.22$ |

So far, we compared sequences without considering possible insertions or deletions in them. Since the best alignment of distantly related protein sequences of ***n*** ≈100 a.a. residues usually requires two or three insertions or deletions of several residues [6–9], these insertions and deletions, together with a shift of one chain relatively to another, totally increase the number ***N*** of independent sequence comparisons by about 3–6 orders of magnitude; thus, the best expected (from Eq. (3)) identity of a ≈100-residue random sequence to “the most similar” to it database protein shifts from the minimal 19-23% to a more realistic 24-31% (see the yellow bar in Fig. 1). For the “half-domain” pieces of the chain this will be shifts even from the minimal 27-33% to a more realistic 34-46% (see Table 1).

Equation (3) and Table 1show that the expected ***M/n*** value decreases with the increasing chain length ***n***. However, one part of a long examined chain can match some part of one protein structure, while another part of this chain can match some part of another protein structure, and AlphaFold 2 is able to do a flexible docking of auch parts (J. Jumper, private communication) using co-evolutionary correlations and restrictions of sequences forming the contacting chain regions (as it does [1, 2] for contacts of remote regions within a continuous chain). This refers to the examined protein chains that cannot be recognized as a whole (and therefore their 3D structures should be considered as “novel folds”): their parts can have a sufficiently high sequence identity to match some parts of different proteins existing in the databases, and AlphaFolds are able to try docking of such parts (J. Jumper, private communication).

## Discussion

With huge UniProtKB and PDB databases currently available, we can conclude that a “new” sequence whose 3D structure is to be identified, typically either has a ≈19-31% sequence identity to some protein whose 3D structure (or that of its close homolog) is already known, or can be divided into a few domain-size parts having the same ≈19-31% sequence identity to some parts of proteins whose 3D structures (or those of their close homologs) is already known, or even into several half-domain-size parts having up to ≈27-46% sequence identity to some parts of proteins whose 3D structures (or those of their close homologs) is already known; and then these parts can be united by a flexible docking. The ≈19-31% sequence identity is typical [4] (Fig. 1) for the members of protein superfamilies whose structural divergence does not exceed 1.7±0.5Å, and ≈27-46% identity is typical for the members of protein superfamilies whose structural divergence does not exceed 1.2±0.3Å (see Fig, 1).

Now we can answer the questions posed at the beginning of this paper. The main reason for the tremendous success of AlphaFolds is (in addition to great and skillful programming) the usage of huge protein databases, which already (as Cyrus Chothia predicted 30 years ago [10]) seem to cover all or almost all of the protein superfamilies existing in nature. Using such databases, multiple sequence alignments and the resulting coevolutionary information (like correlations in pairs [1] and especially in triplets [2] of a.a. residues of the contacting chain regions), AlphaFold *recognizes* a 3D structure framework [11] by a similarity of the examined a.a. sequence to related sequence(s) with already known 3D structure(s), and then only slightly (J. Jumper, private communication) refines the recognized 3D structure using conventional physical potentials and energy minimization.

It is not out of place to mention that the results of the AlphaFold 2 program trained at the databases of 1994 (when the first CASP took place) would be far not as good as now, since databases of 1994 were much smaller: the Swiss-Prot (the ancestor of UniProtKB database) of 1994 only contained about 30000 sequences [12], and the PDB of 1994 about 1000 3D protein structures (https://www.rcsb.org/stats/growth/growth-released-structures). With these databases of 1994, the highest expected (after Eq. (3)) sequence identity for gapless alignments of a 100-residue chain would be not ≈19-23%, as now, but only 15-19% (see the dark-gray bar in Fig. 1), which is below the “twilight zone” [5] where the sequence-based protein structure similarity is not so reliable. As for the alignments with insertions and deletions, we would have, in 1994, more “twilight” recognition with the expected sequence identity of 20-27% (instead of ≈24-31% of now). And it goes without saying that in 1974, when the first international assessment of protein structure prediction [13] took place, and only a dozen of protein 3D structures and a few thousand sequences were known, it would be hardly possible (see a low light-gray bar in Fig. 1) to recognize a protein structure by the AlphaFold programs.

## Conclusions

The main reason for the tremendous success of AlphaFolds is a skillful usage of huge protein databases that have been collected over 60 years. Now they give a possibility to predict, or rather recognize protein spatial structures from their amino acid sequences without considering the process of protein folding [7, 14] that creates these structures.

However, it should be noted that although the assumption that similar sequences have similar folds [4, 7] is practically 100% correct for natural proteins, some specially designed protein sequences demonstrate that there a directed mutation of only one special amino acid residue drastically changes the 3D structure and function [15], which makes the 3D structure recognition from the a.a. sequence in such a designed case rather problematic.

It is also worth noting that AlphaFold 2 contains about 21×10^6^ adjustable parameters (J. Jumper, private communication), which is *at least 1000 times higher* than the number of parameters that are necessary to describe physics of protein chains, including all the pairwise [7], triple, and even quadruple [16] interactions of all atoms existing in proteins (but 21×10^6^ adjustable parameters is close in order of magnitude to the number of quantities needed to describe positions in all rotational and bending degrees of freedom in all 1.5×10^5^ protein structures presented in PDB in 2020).

This 1000-fold excess shows the ratio of AlphaFold’s effort spent on bioinformatics *recognition based on similarity*, and on predictions and refinements based on physics: the former is much greater.

Concluding, I have to emphasize that this paper does not diminish the merit and utility of AlphaFold; it only explains the basis of its success.

## Abbreviations

3D: three-dimensional
a.a.: amino acid
PDB: protein data bank
UniProtKB: a knowledge-based database of protein sequences
Swiss-Prot: a protein sequence data bank
CASP: critical assessment of protein structure prediction meeting
RMSD: a root-mean-square deviation

## Acknowledgements

I am grateful to D.N. Ivankov, N.V. Dovidchenko, S.O. Garbuzynskiy and especially J. Jumper for discussions and E.V. Serebrova for editing the manuscript.

I acknowledge support from the Russian Science Foundation (grant № 21-14-00268).

## References

[1] Senior, A.W., Evans, R., Jumper, J., Kirkpatrick, J., Sifre, L., Green, T., Qin, C., Žídek, A., Nelson, A.W.R., Bridgland, A., Penedones, H., Petersen, S., Simonyan, K., Crossan, S., Kohli, P., Jones, D.T., Silver, D., Kavukcuoglu, K., Hassabis, D. Improved protein structure prediction using potentials from deep learning. Nature 577, 706–710 (2020). https://doi.org/10.1038/s41586-019-1923-7.

[2] Jumper, J., Evans, R., Pritzel, A., Green, T., Figurnov, M., Ronneberger, O., Tunyasuvunakool, K., Bates, R., Žídek, A., Potapenko, A., Bridgland, A., Meyer, C., Kohl, S.A.A., Ballard, A.J., Cowie, A., Romera-Paredes, B., Nikolov, S., Jain, R., Adler, J., Back, T., Petersen, S., Reiman, D., Clancy, E., Zielinski, M., Steinegger, M., Pacholska, M., Berghammer, T., Bodenstein, S., Silver, D., Vinyals, O., Senior, A.W., Kavukcuoglu, K., Kohli, P., Hassabis, D. Highly accurate protein structure prediction with AlphaFold. Nature 596, 583–589 (2021). https://doi.org/10.1038/s41586-021-03819-2.

[3] Callaway, E. What’s next for AlphaFold and the AI protein-folding revolution. Nature 604, 234–238 (2022). https://doi.org/10.1038/d41586-022-00997-5.

[4] Lesk, A.M., Chothia, C. The response of protein structures to amino-acid sequence changes. Phil. Trans. R. Soc. Lond. A 317, 345–356 (1986). https://doi.org/10.1098/rsta.1986.0044.

[5] Chung, S.Y., Subbiah, S. A structural explanation for the twilight zone of protein sequence homology. Structure 14, 1123–1127 (1996). https://doi.org/10.1016/s0969-2126(96)00119-0.

[6] Sunyaev, S.R., Bogopolsky, G.A., Oleynikova, N.V., Vlasov, P.K., Finkelstein, A.V., Roytberg M.A.. From analysis of protein structural alignments toward a novel approach to align protein sequences. Proteins: Structure, Function, and Bioinformatics 54, 569–582 (2004). https://doi.org/10.1002/prot.10503.

[7] Finkelstein, A.V., Ptitsyn, O.B. Protein Physics. A Course of Lectures. 2-nd Edition s, lectures 1-6, 22. Academic Press, An Imprint of Elsevier Science; Amsterdam • Boston • Heidelberg • London • New York • Oxford • Paris • San Diego • San Francisco • Singapore • Sydney • Tokyo (2016). ISBN 9780081012369.

[8] Reva, B.A., Finkelstein, A.V., Skolnick, J. What is the probability of a chance prediction of a protein structure with an RMSD of 6 A? Folding & Design 3, 141–147 (1998). https://doi.org/10.1016/s1359-0278(98)00019-4.

[9] Chan, S.K., Hsing, M., Hormozdiari. F., Cherkasov, A. Relationship between insertion/deletion (indel) frequency of proteins and essentiality. BMC Bioinformatics 8, 227 (2007). https://doi.org/10.1186/1471-2105-8-227.

[10] Chothia, C. One thousand families for the molecular biologist. Nature 357, 543–544 (1992). https://doi.org/10.1038/357543a0.

[11] Roney, J.P., Ovchinnikov, S. State-of-the-art estimation of protein model accuracy using AlphaFold. bioRxiv preprint (2022). https://doi.org/10.1101/2022.03.11.484043.

[12] Bairoch, A., Boeckmann, B. The SWISS-PROT protein sequence data bank and its new supplement TREMBL. Nucleic Acids Res. 22, 3578–3580 (1994). https://doi.org/10.1093/nar/24.1.21.

[13] Schulz, G.E., Barry, C.D., Friedman, J., Chou, P.Y., Fasman, G.D., Finkelstein, A.V., Lim, V.I., Ptitsyn, O.B., Kabat, E.A., Wu, T.T., Levitt, M., Robson, B., Nagano, K. Comparison of predicted and experimentally determined secondary structure of adenilate Kinase. Nature 250, 140–142 (1974). https://doi.org/10.1038/250140a0.

[14] Anfinsen, C.B., Haber, E., Sela, M., White, F. H., Jr. The kinetics of formation of native ribonuclease during oxidation of the reduced polypeptide chain. Proc. Natl. Acad. Sci. USA 47, 1309–1314 (1961). https://doi.org/10.1073/pnas.47.9.1309.

[15] Alexander, P.A., He, Y., Chen, Y., Orban, Y., Bryan, P.N. A minimal sequence code for switching protein structure and function. Proc. Natl. Acad. Sci. USA 106, 21149–21154 (2009). https://doi.org/10.1073/pnas.0906408106.

[16] Pereyaslavets, L.B., Finkelstein, A.V. Development and testing of PFFsol.1, a new polarizable atomic force field for calculation of molecular interactions in implicit water environment. J. Phys. Chem B 116, 4646–4654 (2012). https://doi.org/10.1021/jp212474p.

